# Large-scale identification of protein histidine methylation in human cells

**DOI:** 10.1101/2021.03.31.437816

**Authors:** Sebastian Kapell, Magnus E. Jakobsson

**Author notes:** To whom correspondence may be addressed: Magnus E. Jakobsson.

## Abstract

Methylation can occur on histidine, lysine and arginine residues in proteins and often serves a regulatory function. Histidine methylation has recently attracted notable attention through the discovery of the human histidine methyltransferase enzymes SETD3 and METTL9. There are currently no methods to enrich histidine methylated peptides for mass spectrometry analysis and large-scale analyses of the modification are hitherto absent. In the present study we query ultra-comprehensive proteomic datasets to generate a resource of histidine methylation sites in human cells. We use this resource to explore the frequency, localization, targeted domains, protein types and sequence requirements of histidine methylation and benchmark all analyses to methylation events on lysine and arginine. Our results demonstrate that histidine methylation is widespread in human cells and tissues and that the modification is over-represented in regions of mono-spaced histidine repeats. We also report colocalization of the modification with functionally important phosphorylation sites and disease associated mutations to identify regions of likely regulatory and functional importance. Taken together, we here report a system level analysis of human histidine methylation and our results represent a comprehensive resource enabling targeted studies of individual histidine methylation events.

## INTRODUCTION

Methylation of histidine is a post translational modification (**PTM**) that was first described to occur on actin(1) and myosin(2) proteins around five decades ago. It can occur at two distinct positions denoted as 1-methyl histidine (**1MeH**) and 3-methyl histidine (**3MeH**) (here collectively referred to as **Hme**)(3). Despite being known to the scientific community for long it has gained far less attention than the well-studied protein methylation events on lysine and arginine(3), which are considered as key epigenetic modifications linked to chromatin compaction state and gene activity(4).

Until recently, little was known about the enzymology and significance of Hme. In 2018, SETD3 was uncovered as the first human methyltransferase enzyme targeting histidine and being responsible for the well-established methylation of Actin-H73(5). This finding were shortly thereafter independently validated and functionally shown to modulate smooth muscle contractility(6). In addition, the human METTL18 (also known as C1orf156) enzyme represents a clear homolog to the established yeast methyltransferase Hpm1 (systematic name YIL110W) targeting a RPL3-H243 in *S. cerevisiae*(7, 8), but its enzymatic activity remains to be validated. Finally, human METTL9 has been shown to act as an enzyme with broad specificity generating 1MeH in motifs composed of consecutive histidine residues spaced by small amino acids(9).

Aside from the few recent protein-centric studies focusing on individual Hme events, little is known about the abundance and function of the PTM. PTMs are most frequently studied at a large scale by affinity enrichment of modified peptides, or proteins, followed by mass spectrometry (**MS**)-based identification of targeted sites(10). Such approaches have been described for lysine methylation (**Kme**)(11, 12) and are well-established for arginine methylation (**Rme**)(13). For many PTMs, including Hme, there are no established affinity reagents, creating a need for innovative approaches for characterization. In such cases, querying ultra-comprehensive proteomic datasets for mass shifts corresponding to distinct modification events has recently emerged as a promising alternative(14).

Studies dedicated to global characterization of Hme are until this date absent. To explore the PTM we here mined a panel of ultra-deep human proteome datasets(15) to generate an extensive resource of Hme sites. The analysis revealed that Hme is widespread in human cells, and uncovers its abundance, context, and function in relation to Kme and Rme. To the best of our knowledge, the present study represents the first system level analysis of Hme and is to date the most comprehensive draft of the human histidine methylome.

## MATERIALS AND METHOD

### Querying proteomic hedata for methylation events

Publicly available comprehensive proteomic datasets (ProteomeXchange id: PXD004452) previously published by Bekker-Jensen *et al*(15) were downloaded from ProteomeXchange(16). The datasets were chosen based on exhaustive proteome depth, obtained through extensive off-line peptide fractionation using reverse phase chromatography at alkaline pH and analysis of individual fractions using fast scanning MS methods with a Q-Exactive HF mass spectrometer(15). The analyzed data corresponds to LC-MS/MS analysis of tryptic peptides from human tissue biopsies from colon, liver and prostate as well as the human cell lines A549, HCT116, HEK293, HeLa, MCF7 and SY5Y. In addition, an in-depth analysis of data derived from HeLa cell sample digested with a panel comprised of complementary proteases including trypsin, chymotrypsin, Glu-C and Lys-C was included to achieve comprehensive proteomic coverage.

All raw MS files were searched using MaxQuant(17) (version 1.6.0.17i) against a database containing the canonical isoforms of human proteins (Uniprot Complete proteome: UP_2017_04/Human/UP000005640_9606.fasta) using the default software settings with few exceptions. To reduce the search space, the number of allowed missed cleavages was restricted to one. In addition to the default variable modifications, corresponding to acetylation of protein N-termini and oxidation of methionine, mono-methylation of lysine and arginine, di-methylation of lysine and arginine, tri-methylation of lysine as well as the custom generated modification mono-methylation of histidine were included as variable modifications.

### Bioinformatic analyses

Bioinformatic analysis was performed using the Python programming language: Python Language Reference (version 3.8), available at http://www.python.org. The MaxQuant output files were processed, removing annotated contaminants. Modifications identified in one or more biological replicates containing a mono-, di- or trimethylated modification at an arginine, lysine or histidine were defined as a unique methylation site. If a methylated peptide matched to multiple protein entries, all proteins were categorized as methylated in the downstream analysis. To benchmark our identified sites, the publicly available resource PhosphoSitePlus (version 6.5.9.3)(18) was used as a reference. Protein localization data was derived from the SubCellularBarcode project(19) and the subcellular localization of proteins in the cell line MCF7 was chosen as surrogate dataset in order to infer localization of our identified methylated proteins. Complete predicted subcellular localization of all methylated proteins was also done using the computational algorithm BUSCA(20), allowing for protein assignment into 9 distinct subcellular compartments. Identification of methylations colocalizing with phosphorylation sites were achieved by searching curated phosphoproteomic dataset(21) for known phosphorylations at a distance of 10 amino acids, and the functional score of sites was defined as described in original publication. Annotated protein domains were accessed using the resource portal InterPro (version 82.0)(22) and individual methylation sites was mapped to the interior regions of protein domains annotated in the Pfam database (version 33.1)(23). Functional enrichment analysis was conducted using the String database (version 11.0)(24) and multiple testing was corrected for with the Bonferroni method for false discovery rate (FDR). Logo and enrichment analysis of amino acids flanking methylation sites were performed using the iceLogo server and the human precompiled Swiss-Prot peptide sequence composition as reference(25).

### Proteomic characterization of METTL9 knockout cells

A HAP-1 METTL9 knockout (product number HZGHC004343c010, Horizon Genomics) and a wild type control cell line (product number C631, Horizon Genomics) were propagated and maintained in IMDM Glutamax media (Thermo Fisher Scientific) supplemented with 10% fetal bovine serum (Thermo Fisher Scientific), as well as 100 U/ml penicillin and 100 U/ml streptomycin. The cells were lysed in a guanidine hydrochloride-based buffer and peptides were prepared for analysis using a Q Exactive HF mass spectrometer(26) as previously described(27).

The LC-MS analysis was performed using an EASY-nLC 1200 HPLC system (Thermo Fisher Scientific) coupled to a Q Exactive HF orbitrap instrument. For each single shot proteome analysis, 500 ng peptide was separated using a 3 h chromatography gradient linearly ramped from 10% to 30% buffer B (80% acetonitrile in 0.1% formic acid) in buffer A (0.1% formic acid) during 170 minutes, whereafter the gradient was steeply ramped to 45% buffer B during 10 minutes, before column washing and equilibration. The mass spectrometer was set to continuously sample peptides through a Top12-based data dependent acquisition method. Target values for the full scan mass spectra were set to 3e6 charges in the m/z range 375-1500 and a maximum injection time of 25 ms and a resolution of 60,000 at a m/z of 200. Fragmentation of peptides was performed using higher energy C-trap dissociation (HCD) at a normalized collision energy of 28 eV. Fragment scans were performed at a resolution of 15,000 at a m/z 200 with a AGC target value of 1e5 and a maximum injection time of 22 ms. To avoid repeated sequencing of peptides, a dynamic exclusion window was set to 30 s.

The raw MS files were analyzed using MaxQuant (version 1.6.0.17i) with identical settings to the exploratory searches of published proteomic datasets, statistical analyses was performed using Perseus(28). First, LFQ intensities were imported from the MaxQuant output file denoted “protein groups”. Common contaminants, proteins only identified as modified and proteins hitting the reverse decoy database were thereafter removed by filtering. The resulting data matrix was filtered for proteins detected in at least 70% of the replicates in one experimental condition. The data was then log-transformed and missing values were imputed from the lower tail of the abundance distribution using the default setting in Perseus(28). Proteins displaying significance differences between the conditions were identified through a Student’s T-test (p < 0.05) with P-values corrected for multiple hypothesis testing using the Benjamini–Hochberg method. For cluster analysis, LFQ intensities for proteins displaying a significant difference between the conditions were z-scored and row and columns trees were generated using Euclidean distance and Pearson correlation, respectively. Gene ontology analysis of proteins over- and under-represented in METTL9 knockout cells, was performed using the embedded function in Perseus and P-values were corrected using the Benjamini–Hochberg method.

## RESULTS

Histidine methylation (**Figure 1**) is a poorly characterized PTM which has recently attracted notable attention through the discovery of the human histidine methyltransferase enzymes SETD3(5, 6) and METTL9(9). Large-scale Hme analysis is challenging since there are no available affinity agents to enrich peptides bearing the PTM for MS analysis. Here, we devise an alternative strategy based on mining ultra-deep human proteomic datasets(15) for the modification. This approach enables global identification of cellular Hme events and a subsequent system level analysis of the PTM.

**Figure 1.**
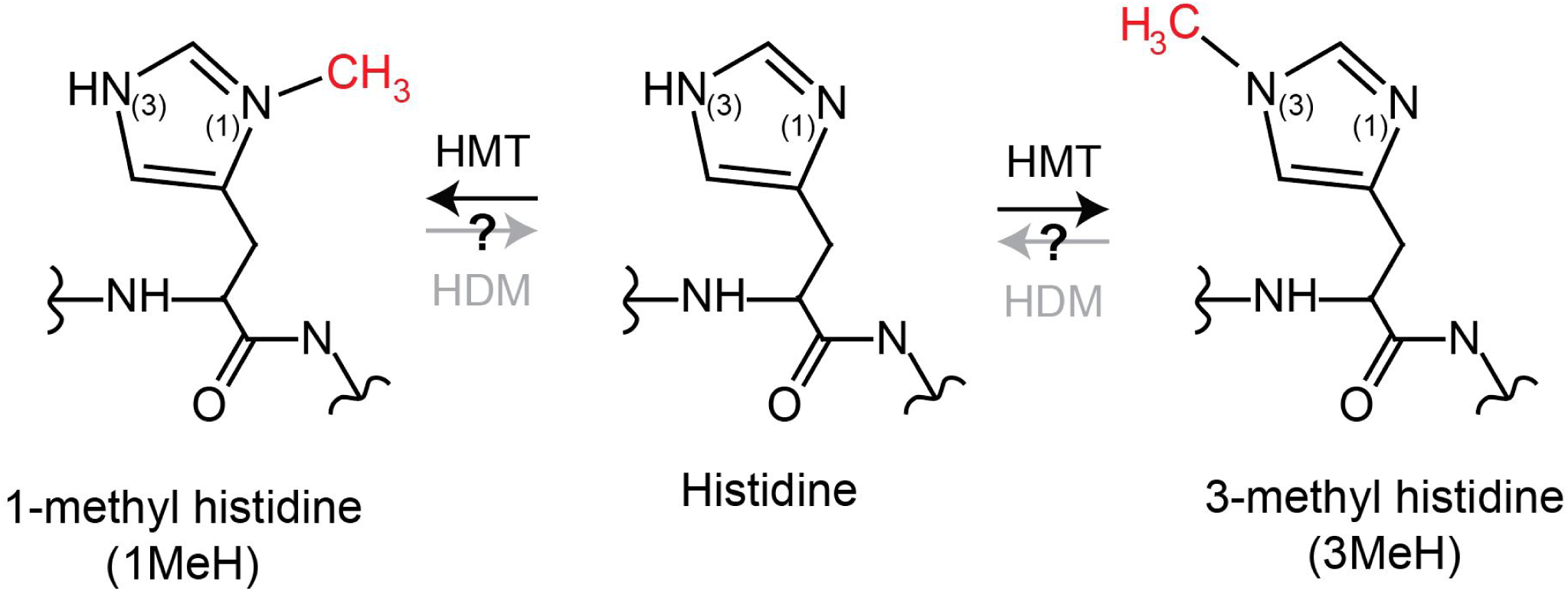
Biochemistry of protein histidine methylation. Structure of the different methylated forms of histidine. The methylations are enzymatically introduced by histidine (<u>H</u>) <u>m</u>ethyltransferases (HMT) and may potentially be removed by histidine (<u>H</u>) <u>d</u>e<u>m</u>ethylases (KDM).

### Histidine methylation is widespread in human cells

To explore the abundance of Hme we searched ultra-comprehensive proteomic datasets derived from the commonly used human cell lines A549, HCT116, HEK293, HeLa, MCF7 and SY5Y as well as tissue biopsies from human colon, liver and prostate(15) (**Figure 2A**). The datasets were selected based on the expression of the recently established human HMT enzymes SETD3 and METTL9 (**Supplementary Figure S1**). The searches were performed using MaxQuant(17) with Hme defined as a custom PTM. To avoid misidentification of Hme sites, the isobaric PTMs mono-, di- and trimethylation of lysine (Kme1, Kme2 and Kme3; referred to as Kme) (**Supplementary Figure S2A**) as well as mono- and di- methylation of arginine (Rme1 and Rme2; referred to as Rme) (**Supplementary Figure S2B**) were defined as additional variable modifications (**Figure 2A**). This approach enables cellular Kme and Rme events to serve as a benchmark for the downstream Hme-centric analyses.

**Figure 2.**
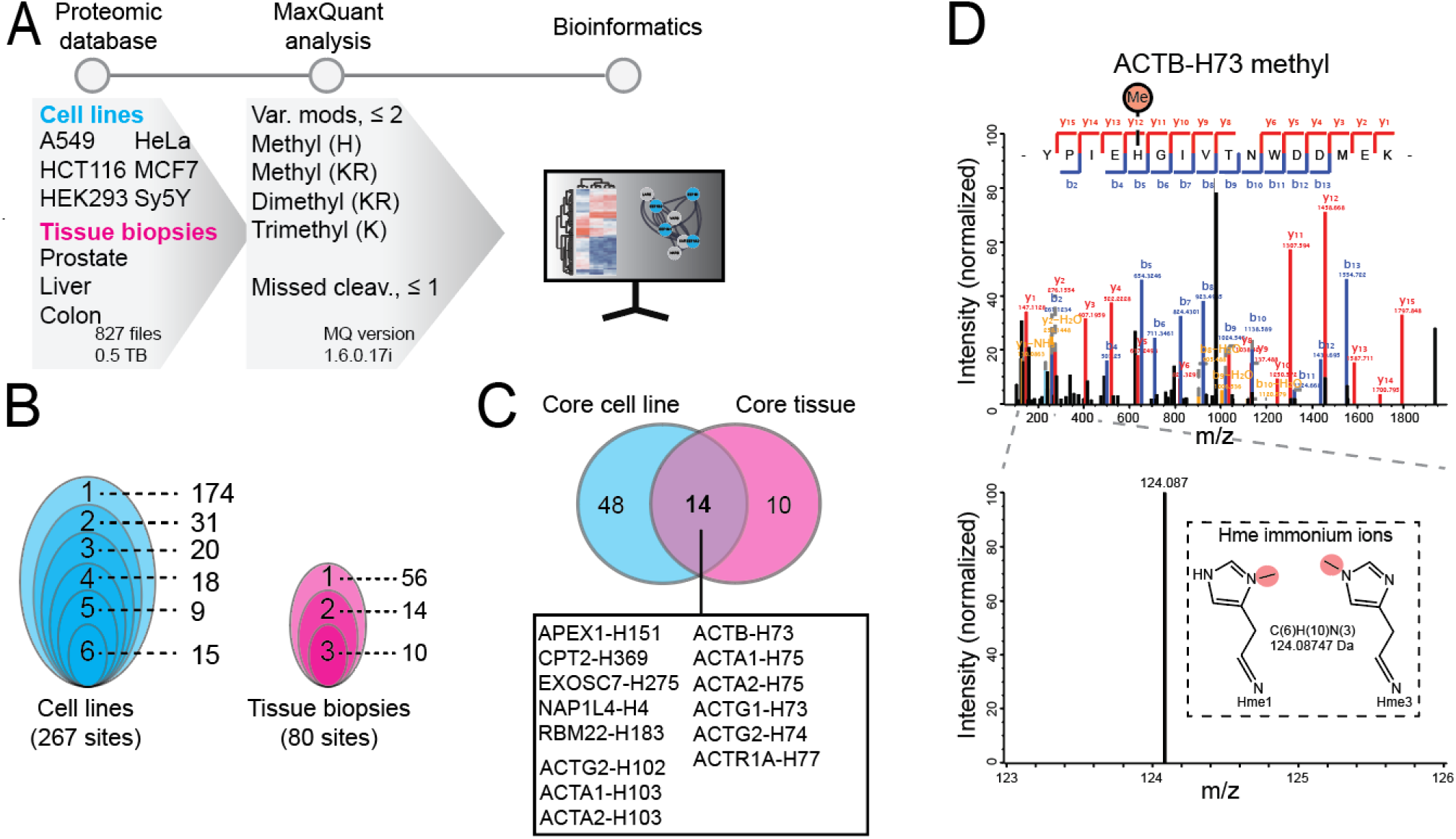
Exploring histidine methylation in human cells and tissues. (**A**) Methylome profiling workflow. Publicly available ultra-comprehensive proteomics datasets from human cell lines and tissue biopsies were searched for histidine, lysine and arginine methylation using MaxQuant. Identified histidine methylation events were analyzed to explore the abundance and cellular context of the PTM. (**B**) Clam plot representation of histidine methylation sites identified in cells and tissues. The total number of sites and the number of shared sites between cell lines (left) and tissue biopsies (right) are indicated. (**C**) Core histidine methylation sites. Sites were categorized as part of the core methylome if identified in more than 50% of the samples in each category. (**D**) Mass spectra supporting methylation of ACTB-H73. Tandem mass for a tryptic peptide covering Y69-K84 of ACTB unambiguously supporting methylation of H73. The presence of a specific immonium ion corresponding to methyl-histidine is indicated.

We anticipated that using a broad range of human cell proteome datasets would enable the identification of both general and cell specific Hme events. The exploratory searches revealed 267 and 80 Hme sites across the cell lines and tissue biopsies, respectively (**Figure 2B**, **Supplementary Table S1**). Moreover, we found several distinct Hme events in multiple cell lines and tissue biopsies (**Figure 2B**). This led us to define a core histidine methylome based on sites identified in 50%, or more, of the cell lines and tissue biopsies (**Figure 2C**). The core methylome includes two sites present in several actin variants, and actin related proteins, corresponding to ACTA1-H103 and established SETD3-target site ACTB-H73(1, 5, 6) (**Figure 2C** and **Supplementary Figure S3A**). Moreover, the core methylome encompasses APEX1-H151 (Uniprot id P27695), CPT2-H369 (P23786), EXOS7-H275 (Q15024), NP1L4-H4 (Q99733) and RBM22-H183 (Q9NW64) (**Figure 2C**) and the Hme sites in these non-actin related proteins do not share apparent sequence homology (**Supplementary Figure S3B**).

A detailed inspection of tandem mass spectra for actin methylation revealed interesting features. First, fragment spectra were of high quality and unambiguously supported methylation of ACTB-H73, with high coverage of up- and down-stream b- and y-ions (**Figure 2D**). Tandem mass spectra from PTM bearing peptides can contain so-called immonium ions with a mass corresponding to the modified residue(29) and their identification may corroborate PTM events(30). Peptides containing unmodified histidine often yield a strong immonium ion with a mass of 110.0718 atomic mass units (**amu**) when fragmented(31). Strikingly, for the tryptic peptide covering ACTB-H73 we instead observed a clear peak at 124.0875 amu, corresponding to the mass of an Hme immonium ion (**Figure 2D**). Similarly, we detected an internal Hme site on the likely histidine methyltransferase METTL18 (METTL18-H154) and the corresponding fragment spectra also contained a clear Hme fingerprint immonium ion (**Supplementary Figure S4**). The observation of METTL18-H154 methylation is interesting as auto-methylation of methyltransferase enzymes is a well-established phenomenon, which has also previously been reported to occur with an amino acid specificity reflecting the enzymes physiological substrate(7). In summary, the above analysis demonstrates that cellular Hme sites can be identified by querying comprehensive proteomic datasets and that the PTM is widespread in human cells and tissues.

### Exploring the HeLa methylome

Intrigued by our observation that Hme is prevalent in human cells we embarked on an in-depth exploratory analysis of the PTM. For this analysis we focused on human HeLa cells, which is arguably the most used human cell line model in experimental research. We devised the now established successful approach of querying publicly available comprehensive datasets for Hme. This time, the chosen datasets(15) were specific for Hela cells and generated using a panel of complementary proteases including, chymotrypsin, Glu-C and Lys-C, aside from trypsin, allowing for unprecedented proteome depth and coverage(32).

Across the different proteases, the analysis revealed support for 2526 distinct cellular methylation events at 2241 sites (**Figure 3B**). The number of methylation events exceed the number of sites since several individual arginine and lysine residues were detected with varying degrees of methylation (**Supplementary Table S1**). Roughly 12% of methylation events correspond to Hme (n=299) and the modification was less prevalent (n=299) than Kme (n=895) and Rme (n=1332) (**Figure 3B**). In line with these observations, a previous study has suggested that roughly 14% of protein methylation events occurs on histidine(33).

**Figure 3.**
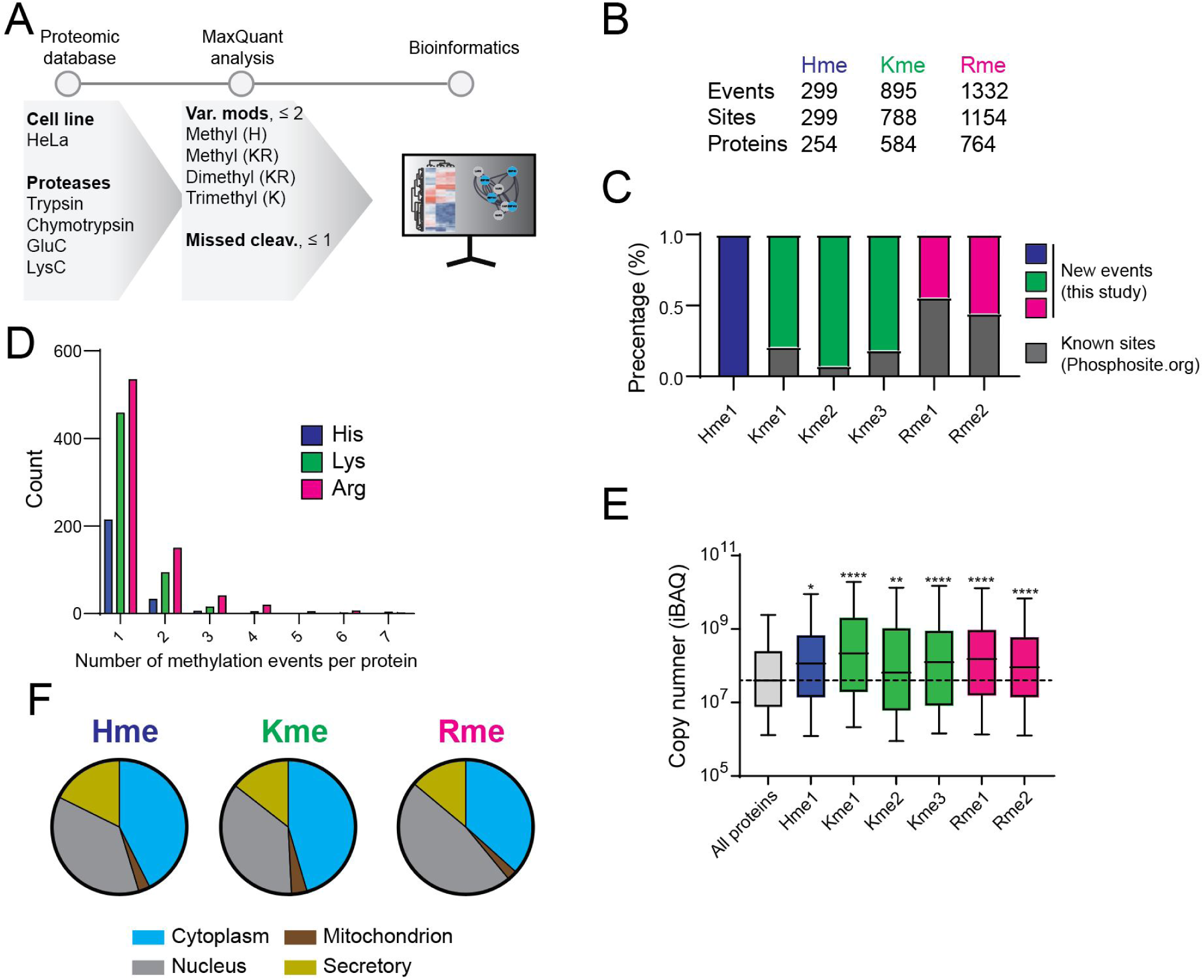
In-depth characterization of the HeLa methylome. (**A**) HeLa-specific methylome profiling workflow. A panel of comprehensive proteomic datasets generated using several proteases to obtain extensive proteome and PTM coverage were searched for histidine, lysine and arginine methylation. The identified histidine methylation events were explored using a range of bioinformatic tools and benchmarked to identified lysine and arginine methylation. (**B**) Total counts of distinct methylation events, methylation sites and targeted proteins are indicated. (**C**) The percentage of sites identified in this study as compared to the dataset available from PhosphoSitePlus. (**D**) The number of methylation events per protein. (**E**) Cellular abundance of methyl proteins. The distribution of iBAQ values for proteins harboring a methylation site is shown. Significance was assessed for each group compared to control (All proteins) by multiple comparison using one-way ANOVA, (adjusted P value <0.01). (**F**) Subcellular localization of proteins for Hme, Kme, Rme assigned to a designated compartment neighborhood as described in the SubCellBarcode project. Each methylation type as a relative distribution in the nucleus, cytoplasm, mitochondria and secretory compartment.

A comparative analysis of our HeLa methylome data to publicly available PTM resources(18) suggests that the bulk of (>80%) of Hme and Kme sites are not previously characterized (**Figure 3C**). The fraction of novel sites was notably lower for Rme (**Figure 3C**), which can be expected as established workflows for affinity enrichment and MS characterization of the PTM exist(13). The large number of identified new Hme sites highlights that the generated dataset is suitable for an exploratory systems level analysis of the PTM.

To investigate whether Hme is scattered across the proteome, or frequently occurring on individual proteins, we first analyzed the number of Hme events per Hme protein. This analysis revealed that a single methylation event was identified for most (>80%) Hme proteins and that no proteins were identified with more than two Hme sites, and similar trends were observed for both Kme and Rme (**Figure 3D**). Next, we explored the abundance of identified methyl proteins. Interestingly, we found that Hme, Kme and Rme were all overrepresented on abundant proteins (**Figure 3E**) and envision two alternative explanations for this finding. First, the proteomics datasets in this study were generated using data dependent acquisition MS, an approach intrinsically biased to identify abundant peptides and PTMs(34). Alternatively, it has been suggested that certain methyltransferases have evolved specificity towards key abundant cellular proteins to modulate key cellular functions(35). Prominent examples include Kme and Rme in the core histone H3(36), the key translational elongation factor eEF1A(37, 38), and the molecular chaperone Hsp70(39–41) as well as Hme in actin(1, 5, 6), targets that were all identified and validated in this study (**Supplementary Table S1**).

Next, we investigated the subcellular localization of methyl proteins. To this end, we used both experimental data based on the SubCellBarCode resource(19) as well as the predicted localization based on the BUSCA approach(20). Both methods place Hme, Kme and Rme proteins in the cytoplasm, nucleus, mitochondria and in secretory compartments, at comparable frequencies (**Figure 3F** and **Supplementary Figure S5**), suggesting that Hme, Kme and Rme are all widespread across cellular structures.

In summary, the above indicates that Hme is prevalent and present in all major cellular compartments.

### Domains and motifs targeted by Hme

Intrigued by the finding of Hme events across different cellular compartments, we next used the Pfam(23, 42) database to explore which proteins families and domains that are targeted. Although Hme, Kme and Rme proteins displayed similar subcellular localization profiles (**Figure 3F**) they are clearly enriched in different Pfam entries (**Figure 4A**). Reassuringly, the most strongly enriched Pfam entries include to domains where the methylations are well established to exert important function including Actin for Hme(5, 6), RNA recognition for Rme(13) and core histone proteins for Kme(4) (**Figure 4A**). Notably, Hme was found overrepresented in Pfam entries associated with zinc binding properties (E3 Ligase, CCCH-zinc finger; Zinc finger C2H2 type; Zinc finger CCCH type; Zinc finger C-x-C-x-C type (and similar); ZIP zinc transporter) (**Figure 4A** and **Supplementary Figure S6**).

**Figure 4.**
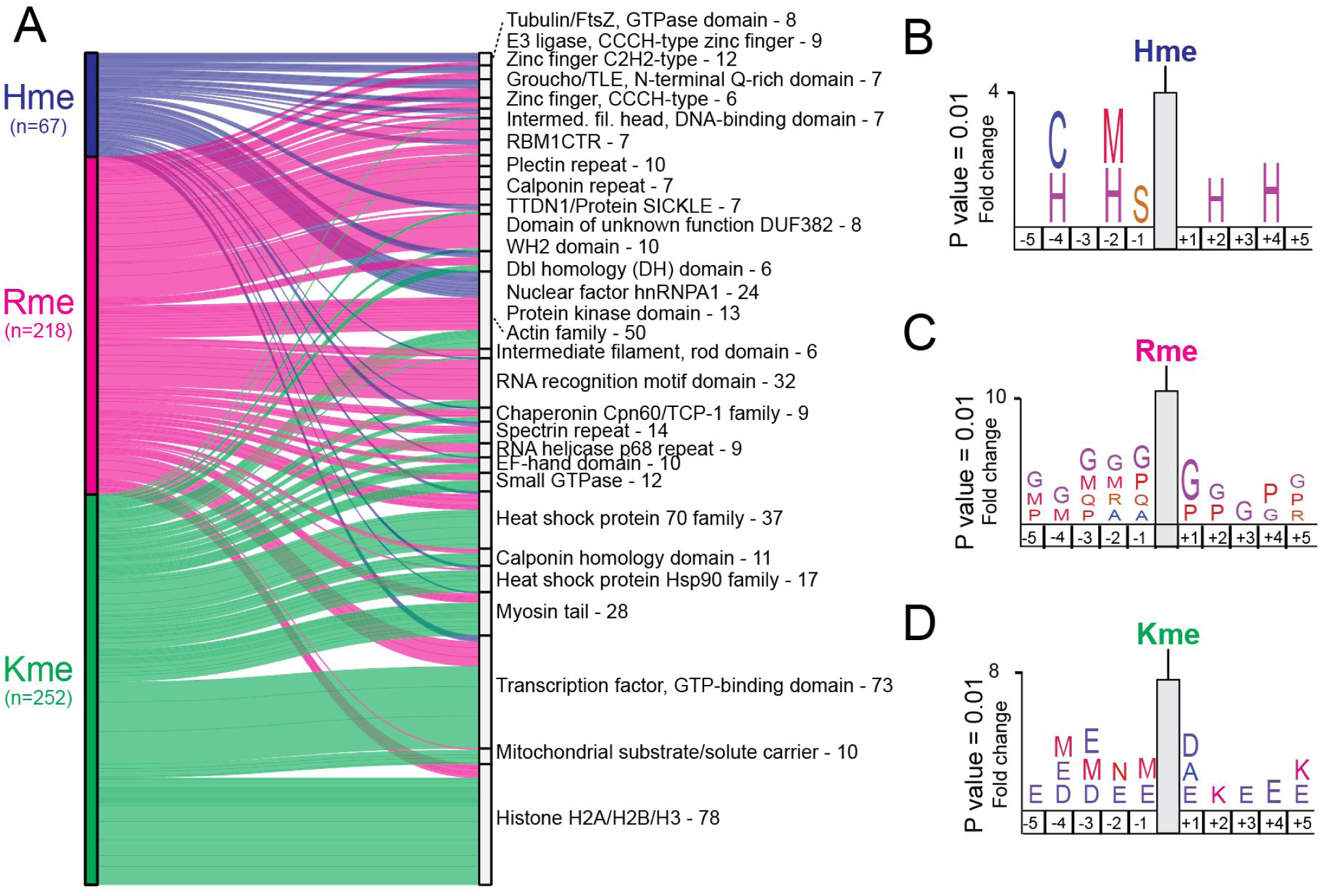
Domains and sites targeted by methylation. (**A**) Sankey plot illustrating the top 30 Pfam domains targeted by histidine, arginine and lysine methylation in HeLa cells. Total number of unique methylation sites residing within an annotated protein domain in the Pfam database is indicated. Statistical analysis for significantly enriched domains is shown in **Supplementary Figure S6**. (**B-D**) Methylome sequence logos. Logos illustrating over representation of amino acids in the 5 positions up- and down-stream of identified (**B**) Hme, (**C**) Rme, and (**D**) Kme sites. Full sequence logos and heatmap analysis are shown in **Supplementary Figure S7**.

Having established that Hme is over-represented in certain zinc binding proteins and domains, we next set out to analyze the sequence context flanking Hme sites using the iceLogo approach(25). We queried the 5 flanking amino acids for all detected Hme, Kme and Rme sites using the human precompiled Swiss-Prot peptide sequence composition as reference. This analysis revealed distinct sequence preferences for the different methylations (**Figure 4B-D**). Hme was overrepresented in mono-spaced repetitive histidine (H) sequences (**Figure 4B**), Rme in glycine (G) rich regions (**Figure 4C**), and Kme in the context of the acidic residues aspartate (D) and glutamate (E) (**Figure 4D**).

In summary, the above results demonstrate that Hme is over-represented in specific classes of zinc binding proteins and in the primary sequency context of consecutive mono-spaced histidine residues.

### Co-occurrence of methylation with phosphorylation and disease associated mutations

It has been reported that methylation events can cross-talk with other PTM types such as phosphorylation, constructing a regulatory circuit known as methyl-phospho switch(43). For example, the lysine methyltransferase SET7 have been shown to regulate the stability of DNA methyltransferase-1 (DNMT1)(44), a key enzyme in maintaining methylation patterns after DNA replication. DNMT1 is methylated at Lys142 and the adjacent Ser143 which can be phosphorylated by AKT1 kinase. These two PTMs are mutually exclusive and in absence of phosphorylation at Ser143, DNMT1 is methylated at Lys142. The consequence of the methylation is an overall decrease in abundance of the key epigenetic regulator DNMT1. In order to pinpoint specific Hme events of high functional importance we integrated a highly curated phosphoproteomic dataset into our analysis(21). We searched for phosphorylations on serine, threonine or tyrosine within a distance of 10 amino acids from a methylation event and this analysis revealed 1999 co-localizing phosphorylation sites (**Supplemental Table S2**). A distinct advantage of the curated phosphoproteomic dataset was that it had been evaluated using a novel machine learning model that integrated multiple features relating to conservation and structural properties of the phosphorylation site, that are indicative of functional relevance, thus providing a functional score. This allowed us to evaluate our identified methylation events based on the functional score of the phosphorylation site in close proximity (**Figure 5A**). We found that Hme, in addition to Kme and Rme, co-localized with phosphorylation sites with an above median functional score (**Figure 5A**).

**Figure 5.**
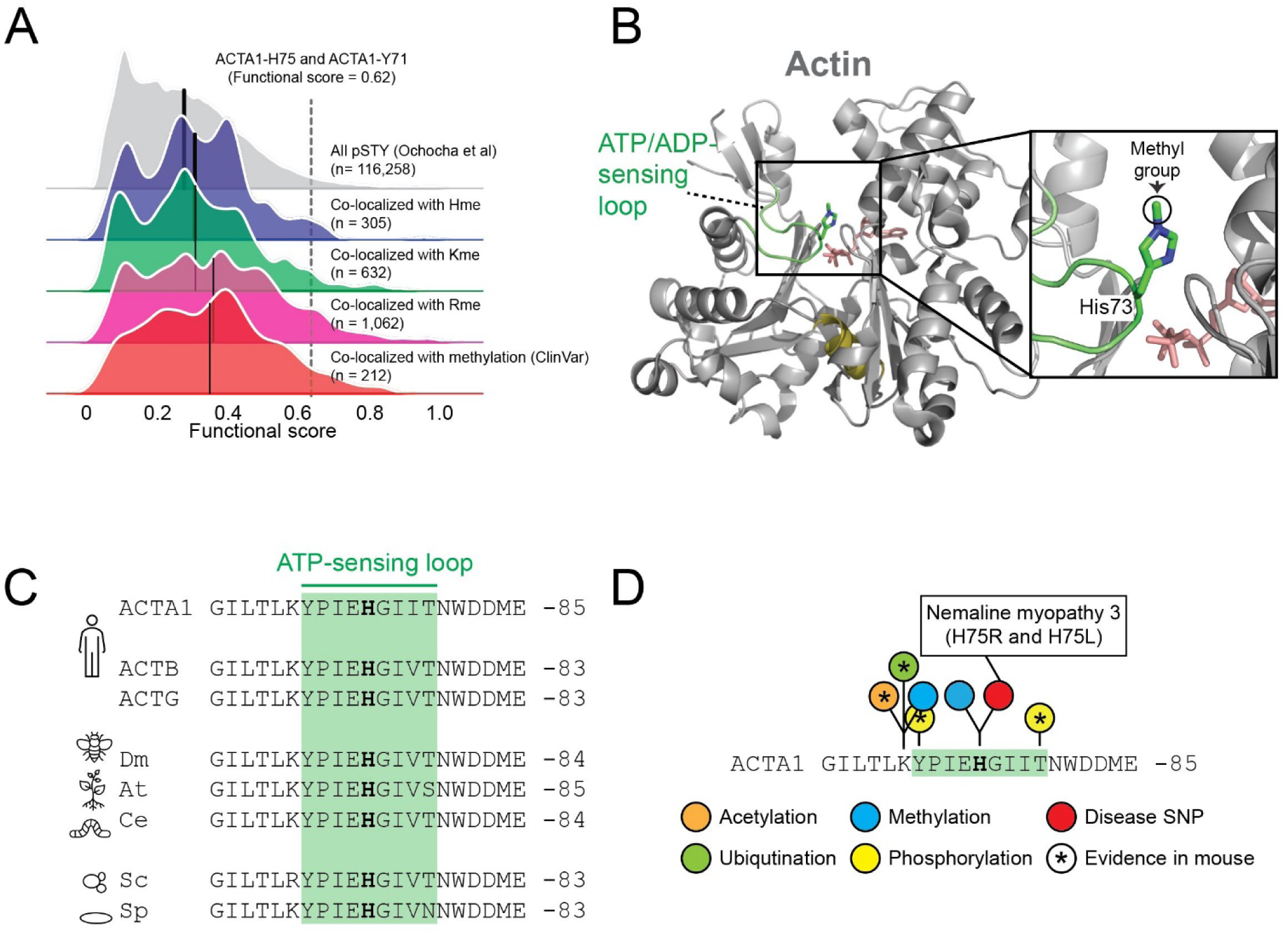
PTM colocalization. (**A**) Kernel density plots for the functional score distributions of colocalizing phosphorylation sites with identified methylation events. Subsets of phosphorylation sites colocalizing with Rme, Kme or Hme methylation sites plotted separately. Separate grouping of phosphorylation sites co-localizing with methylation sites with a reported polymorphism or mutation associated with a pathological condition (ClinVar) is shown. Black line indicate group mean. Methylation of ACTA1-H75 and the colocalizing phosphorylation event on ACTA1-Y71 are indicated. (**B**) The structural context of actin histidine methylation. The structure of actin is shown in cartoon representation whereas ATP and the methylated histidine residue H73 is shown in stick representation. The hinge region (olive), ATP (salmon) and the H73 containing ATP-sensing loop (green) are indicated. The structure is derived from rabbit muscle alpha actin (PDB #1EQYE). (**C**) Evolutionary conservation of the methyl-histidine containing loop in actin. Sequences: human ACTA1 (P68133), ACTB (P60709) and ACTG (P63261) as well as homologues from Drosophila melanogaster (dm; AAA28314.1), Arabidopsis thaliana (at; NP_187818.1), Caenorhabditis elegans (ce; NP_508841.1), Saccharomyces cerevisiae (cs; NP_116614.1) and Saccharomyces pombe (sp; NP_595618.1). (**D**) The methyl-histidine containing loop in ACTA1 is a PTM hotspot. PTMs annotated in the PhosphoSitePlus database (v6.5.9.3) are shown. Modifications observed in mouse ACTA1 (star) and sites corresponding to disease associate mutations are indicated (red).

In addition, we queried a publicly available database(45) for mutations in amino acid positions undergoing a methylation event in order to identify sites linked to pathological conditions(46). This analysis uncovered mutations of 212 methylation sites that are co-localizing with phosphorylations (**Supplemental Table S3**). Notably, the functional score of phosphorylation sites colocalizing with methylation site mutations also had an above the median functional score (**Figure 5A**).

Our analysis suggest that protein methylation co-localizes with functionally important phosphorylation sites, suggesting crosstalk between the PTMs. However, whether crosstalk has a relevance in relation to regulation of cellular function needs to be further experimentally validated for each individual protein.

### Identification of a conserved PTM hotspot in actin

Intrigued by the observations that the ACTA1-H75 methylation site colocalizes with the functional high-scoring ACTA1-Y71 phosphorylation site that is mutated in severe nemaline myopathy(47) as well as the phosphorylation of the adjacent ACTA1-Y71 site (**Figure 5A** and **Supplemental Table S2**), we decided to do an in depth analysis of the region. Structurally, ACTA1-H75 is located in a loop which has been reported to sense nucleotide binding(48) (**Figure 5B**). The loop is perfectly conserved between humans, fly, plant, worm and yeast, emphasizing its functional importance (**Figure 5C**). To explore whether the ATP-sensing loop is targeted by additional PTMs, we again queried the PhosphositePlus database. This analysis revealed five annotated PTMs in the H75-containing nucleotide sensing loop and its flanking residues (**Figure 5D**), corresponding to mono-methylation, ubiquitination, and acetylation of K70 as well as phosphorylation of Y71 and T79. Notably, the bulk of these PTMs have also been observed in mouse actin (**Figure 5D**), indicating that modification of this loop is evolutionary conserved.

In summary, we observed disease associated mutations and PTMs at, and in proximity to, ACTA1-H75, a Hme site previously categorized as belonging to the core Hme-ome. The findings emphasize the functional importance of the ATP-sensing loop harboring H75 and support a model where multiple PTMs may play a role in actin regulation.

### Cellular effects of METTL9 KO

Having uncovered hundreds of cellular Hme events and explored the subcellular distribution and context of the PTM, we sought to also investigate its direct biochemical functions. Human METTL9 was very recently described as a histidine methyltransferase introducing the bulk of 1MeH in mammalian proteomes through methylation of histidine in the context of a HxH motif, where “x” is a preferentially small amino acid such as alanine, glycine or serine(9). The reported specificity of METTL9 corresponds perfectly to our observed over-represented motif for Hme (**Figure 4B**), highlighting METTL9 gene targeted cells as a suitable model for studies of cellular Hme loss.

To uncover cellular processes regulated by Hme, we therefore obtained a CRISPR-mediated METTL9 knockout (KO) HAP-1 cell line(49) and characterized its steady-state proteome, compared to a wild type (WT) control (**Figure 6A**). Cellular proteins were extracted and quantified using a label free MS approach (**Figure 6A**). Principal component analysis of protein intensities revealed a clear separation of WT and KO cells in the first component (**Figure 6B**), indicating that the difference between the cellular conditions exceed the technical experimental variation. Accordingly, hierarchical cluster analysis of proteins with a significant difference (Student’s T-test, P-adj < 0.05) in abundance between the conditions revealed two distinct clusters of over- and under-represented proteins in the KO cells (**Figure 6C**). To obtain insights into the processes and functions affected in the METTL9 KO cells we performed gene ontology analyses of the clusters. This analysis revealed vesicle and vesicle-related processes as under-represented and nuclear nucleic acid-associated processes as over-represented in the KO cells (**Figure 6D**). This suggests a complex and pleiotropic phenotype caused by METTL9 deletion and loss of pervasive Hme1.

**Figure 6.**
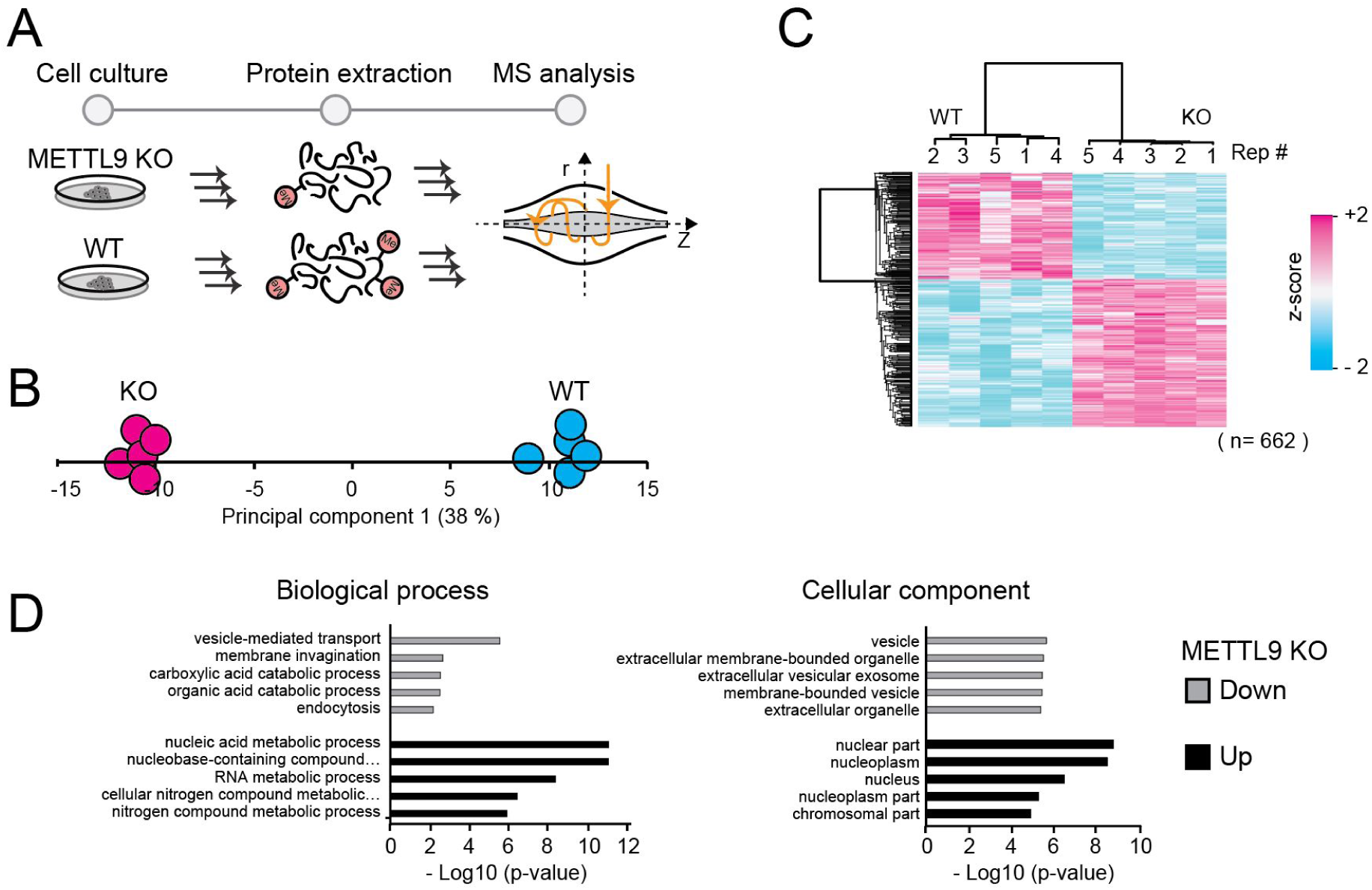
Cellular effects of METTL9 knockout. (**A**) Workflow for proteomics characterization of METTL9 KO cells. Proteins were extracted from HAP-1 wild type (WT) and METTL9 knockout (KO) cells and processed for label free quantitative mass spectrometry. (**B**) Principal component analysis. Separation of experimental conditions (WT and KO) and replicates (n=5) in the first principal component is shown. (**C**) Clustering analysis. Hierarchical cluster of z-scored LFQ intensities for proteins having a significant difference in abundance (Student’s T-test, P-adj < 0.05; Benjamini-Hochberg FDR) between the WT and KO condition. (**D**) Gene ontology analysis. The top five ontologies within biological process and cellular component are shown both for up- and down regulated proteins in METTL9 KO cells, relative to a WT control.

## Discussion

We here report the first large-scale analysis of Hme and demonstrate that the PTM is widespread in human cells and tissues. Our analyses indicate that the PTM is present in all major cellular compartments and that it is overrepresented in specific protein families, in particular in actin and in zinc-binding proteins. Taken together, we present the hitherto most extensive resource on cellular Hme events and perform the first system level analysis of the PTM.

Global PTM analysis almost invariably involves PTM affinity enrichment followed by MS analysis. Such approaches have contributed greatly to the knowledge on phosphorylation(50, 51), acetylation(52), ubiquitination(53), SUMOylation(54) and Rme(13) but it has been more challenging to generate robust affinity agents for proteomics characterization of Kme, and to the best of our knowledge, has not yet been tried for Hme. To study Hme, we here deployed an alternative brute-force approach, taking advantage of the high throughput of modern mass spectrometers, and queried ultra-comprehensive proteomic datasets for the PTM. To provide a benchmark for identified Hme events, we also searched the datasets for Rme and Kme. The strength of our approach can be highlighted by comparing our identified Kme-ome to the current state-of-the-art Kme proteomics studies. We identified 895 Kme events in HeLa cells alone (**Figure 2B**). For comparison, a study Cao et al(12) using lab-specific IgGs for all Kme states identified 552 Kme events in HeLa cells by another acknowledged study by Moore et al(11) using the bispecific Kme1 and Kme2 binding 3xMBT domain revealed 31 Kme events in 293T cells.

A drawback with our approach is the extensive laboratory work associated with off-line fractionation of peptides before MS analysis required to generate ultra-comprehensive proteomic datasets. Moreover, searching comprehensive datasets for several variable PTMs is both computationally challenging and time consuming. The establishment of Hme affinity enrichment workflows for MS would reduce the requirements for MS analysis time and data processing, and drastically reduce the efforts and costs associated with our Hme-omic approach. Thus, we foresee efforts will be made towards the generation of Hme-specific affinity agents and their optimization for Hme-proteomic applications. The affinity agents can be generated though classical animal immunizations but this approach is inherently associated with low reproducibility(55). A more robust and reproducible approach would be to generate recombinant Hme-binders, using phage-displayed recombinant antibody libraries, an approach proven feasible for the PTM sulpho-tyrosine(56).

Affinity enrichment followed by MS is arguably the most widely employed approach to study PTMs. A prominent example of this is for phosphorylations, where affinity agents that specifically bind to the modified functional phosphate group exist(51). These rely on immobilized metal cations with affinity for the negatively charged phosphate group and can consequently be used for enrichment of phosphorylated serine, threonine and tyrosine(10). For protein methylations, which are small and chemically subtle PTMs, there are no such affinity agents available. Antibodies and specific methyl-state binding protein domains have instead been used to enrich Kme and Rme modified peptides for MS(11–13). However, antibodies and domains often display a preference for PTMs in specific contexts. For example, several Rme antibodies have a preference for flanking residues(57) and the affinity of Kme-binding domains can be affected by neighboring PTMs, such as phosphorylation(58). Although the depth of the histidine methylome may conceivably be increased beyond this study through future affinity enrichment-based approaches, the herein identified Hme events, and the subsequent bioinformatic analysis, is not skewed or biased through context-specific affinity agents.

Our proteomic characterization of METTL9 KO cells suggests a complex phenotype with perturbations of both vesicle-associated and nuclear-linked cellular process. A recent study suggested the presence of as many as 2807 candidate METTL9 target sites (HxH, x = A, N, G, S or T) in the human proteome(9). Our observed pleiotropic phonotype for METTL9 KO cells may be linked to the large number of plausible substrates for the enzyme, which will likely be subjects for future studies. The large number of substrates for METTL9 also render our METTL9 targeted cells a poor tool to study biochemical functions of individual Hme events. Instead, we suggest ectopic expression of WT and HxH-mutated METTL9 substrates in cells as a preferred approach.

Actin proteins are subject to a wide range of PTMs, many of which have determined regulatory function(59). One prominent example is SETD3 mediated methylation of ACTB-H73, which corresponds to ACTA1-H75, which modulates actin dynamics by accelerating the assembly of actin filaments, a process preventing primary dystocia(6). Another example, is a unique multi-step N-terminal processing and modification machinery involving N-terminal acetylation of the initiating methionine (iMet), followed by excision of iMet and subsequent acetylation of the residue in position 2(60). Our integrated analysis of Hme colocalization with machine learning predicted functional importance scored phosphorylation events (**Figure 5A**) uncovered ACTA1-H75 as colocalizing with ACTA1-Y71 phosphorylation, a site attributed with a high function score (0.62)^20^. Interestingly, we observed five additional PTMs in the six up- and downstream residues, of these four had also been reported to occur in mouse (**Figure 5D**). To the best of our knowledge, so far only H75 methylation has been the subject of detailed biochemical studies. The multiple modifications within the loop reassembles the numerous PTMs in the flexible tail of histone H3, a key component of the histone code determining chromatin compaction state and gene activity. Given the extent of PTMs in the functionally important ATP-sensing loop, a similar “actin code” may exist, where multiple dynamic PTMs collectively, or individually, determine the molecular functions of actin.

The protein methylation dataset we have generated may support further studies on Hme and we envision three direct applications. First, synthetic peptides corresponding to the relatively small HeLa Hme-ome (n=299) can be generated and evaluated as substrates for new candidate histidine methyltransferase enzymes. The human genome encodes over 200 predicted methyltransferases(61) and given the abundance and sequence diversity of identified Hme events (**Supplementary Table S1**), a considerable fraction of these may catalyze Hme. Candidate histidine methyltransferases may be cloned, expressed, and isolated from bacterial systems, and evaluated for activity on peptide arrays comprising the Hme-ome. Second, a synthetic peptide library corresponding to the Hme-ome may be used to uncover Hme-driven protein interactions for yet not discovered Hme reader proteins and Hme demetylases through affinity-enrichment MS approaches(27, 62). Third, our resource provides the necessary information to design large-scale targeted MS methods(63) for Hme that can be used to further explore the regulation and variation of Hme-ome in human cells, tissues, and biological fluids.

In summary, our data extends the current knowledge of Hme and the study represents the first system level analysis of the PTM. Finally, we encourage the research community to use this resource for large-scale targeted MS and detailed biochemical studies of individual sites to shed further light on the emerging field of protein methylation biology.

## DATA AVAILABILITY

Available on request to

## ACKNOWLEDGEMENTS

We would like to thank the Proteoforms@LU mass spectrometry infrastructure research team for useful discussion and manuscript input, Jesper V. Olsen for providing early access to proteomics datasets and Júlia Szántó for assistance with HAP-1 cell culture.

---

**Supplementary Figure S1.**
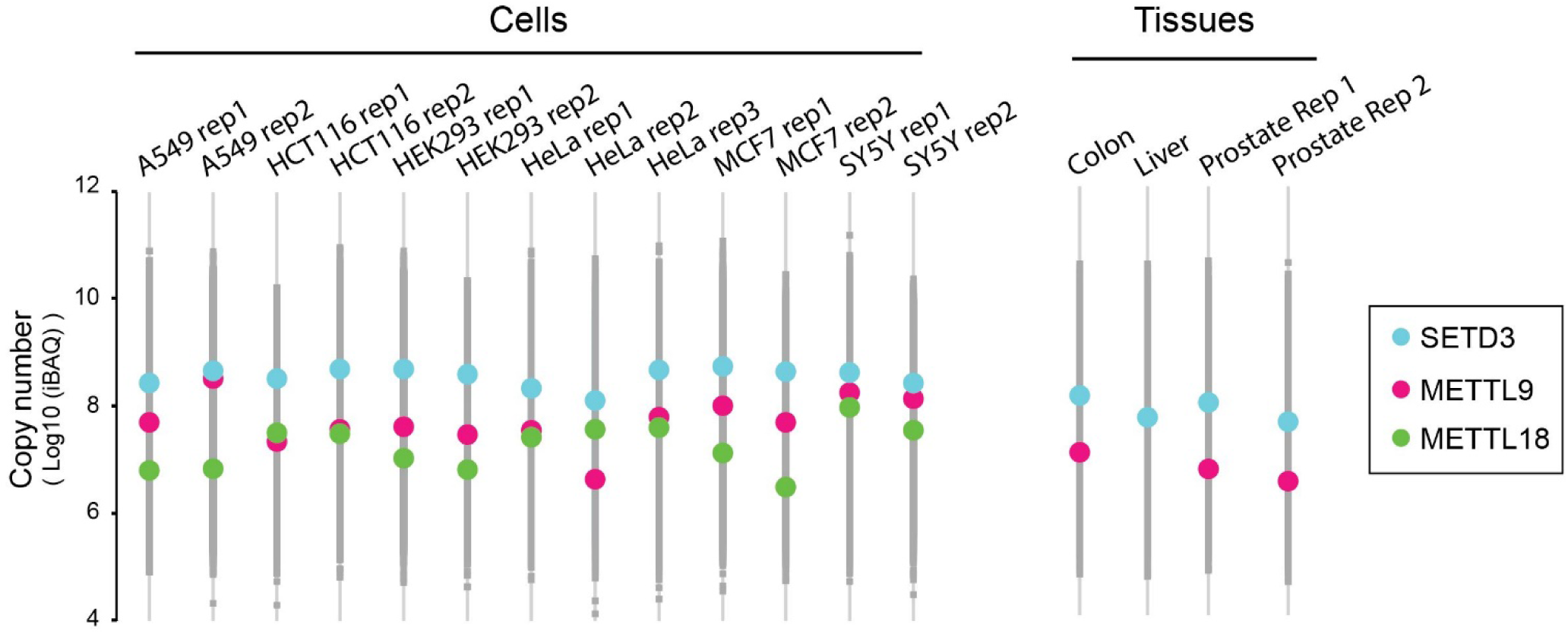
Expression of histidine methyltransferases in cells and tissues. Profile plot showing the cellular copy number (iBAQ value) for the established human histidine methyltransferase enzymes SETD3 and METTL9 as well as the candidate histidine methyltransferase METTL18.

**Supplementary Figure S2.**
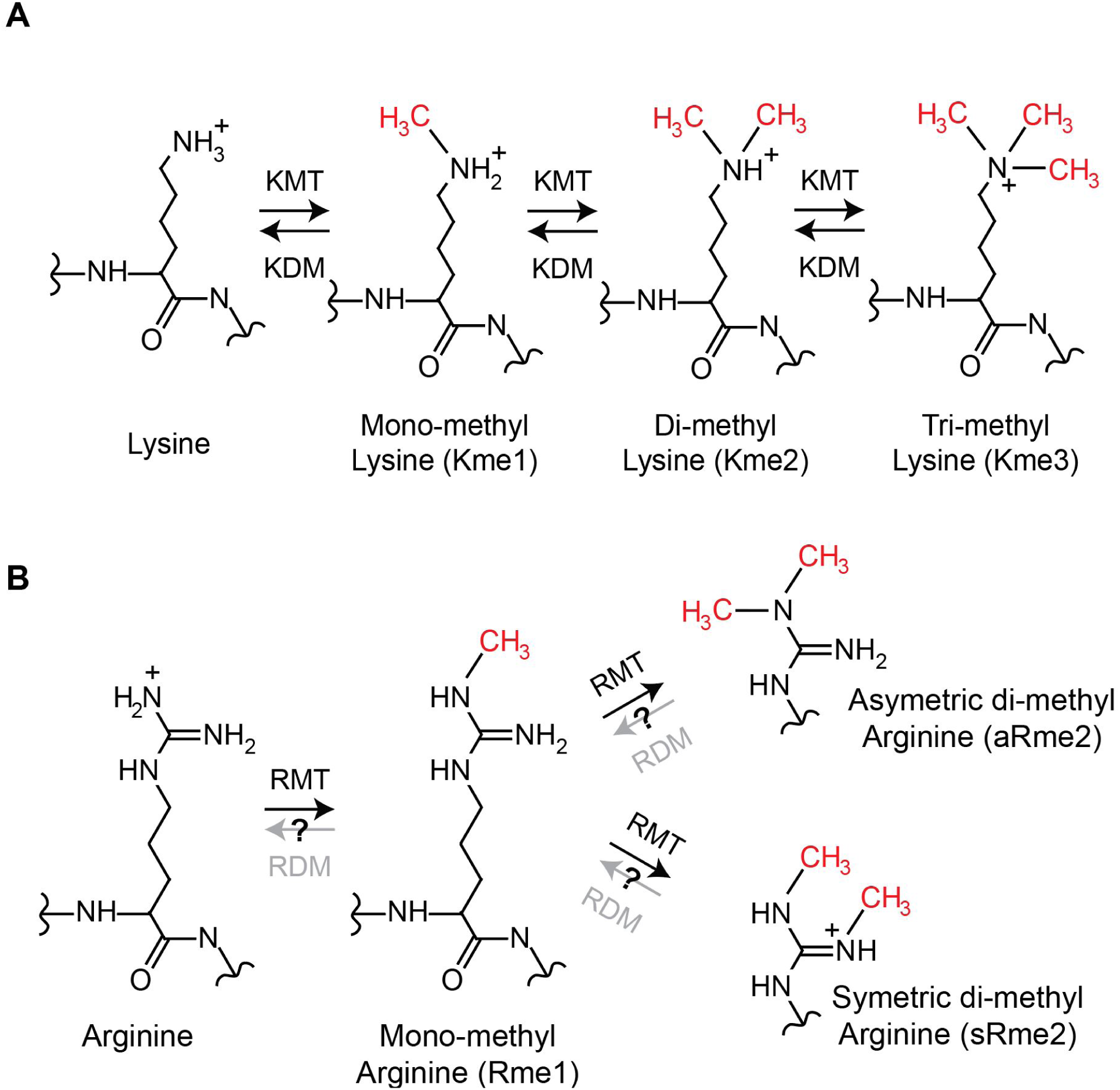
Biochemistry of lysine and arginine methylation. (**A-B**) The chemical structures and enzymology for (**A**) lysine methylation and (**B**) arginine methylation are shown. The methylations are introduced by lysine methyltransferase (KMT) and arginine methyltransferase (RMT) enzymes, respectively. Lysine methylation can be reversed by lysine demethylases (KDM). It is hypothesized, but not yet established that arginine demethylase (RDM) enzymes exist.

**Supplementary Figure S3.**
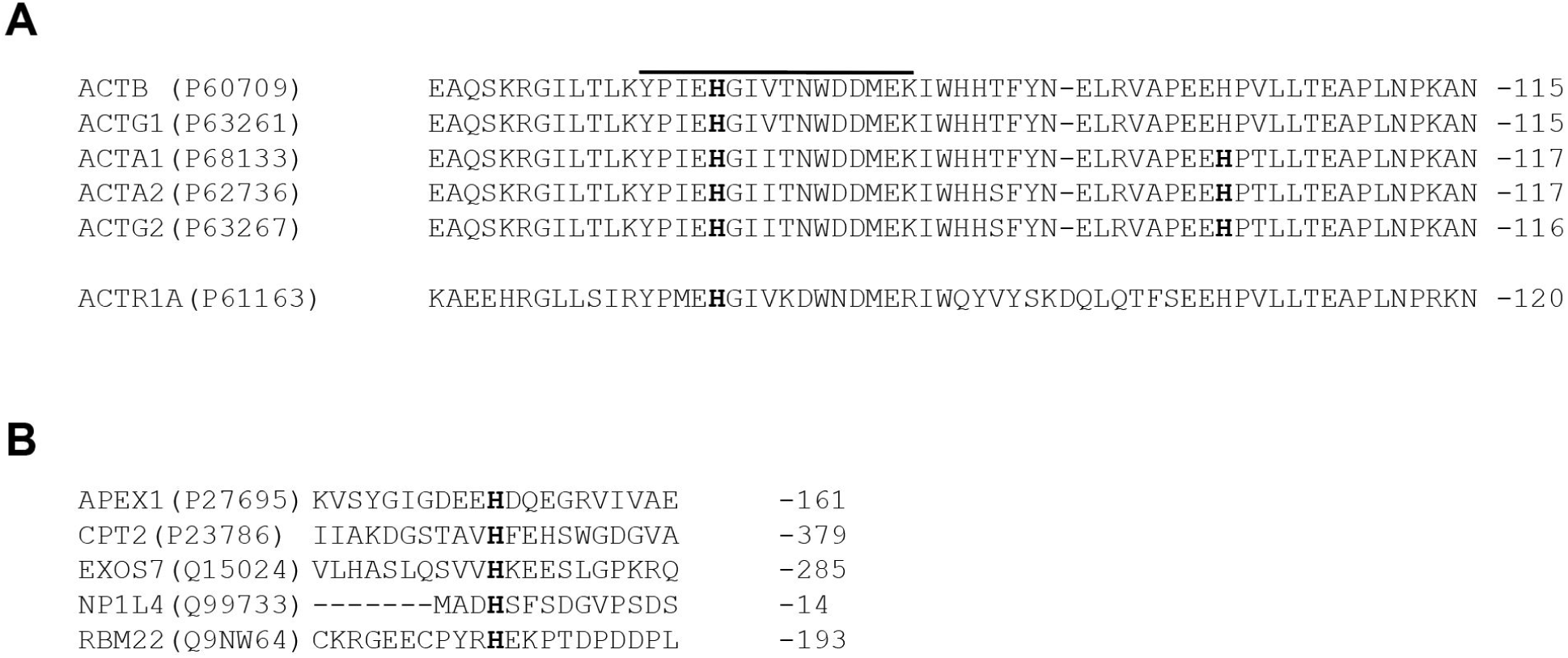
Sequence alignments of core histidine methylome proteins. (**A**) Protein sequence alignment of human actin proteins. Sites of which methylation is supported in our analysis are indicated in bold. (**B**) Protein sequence alignment of non-actin core histidine methylome proteins. Proteins shown are APEX1-H151 (Uniprot id P27695), CPT2-H369 (P23786), EXOS7-H275 (Q15024), NP1L4-H4 (Q99733) and RBM22-H183 (Q9NW64). The histidine detected as methylated is highlighted in bold.

**Supplementary Figure S4.**
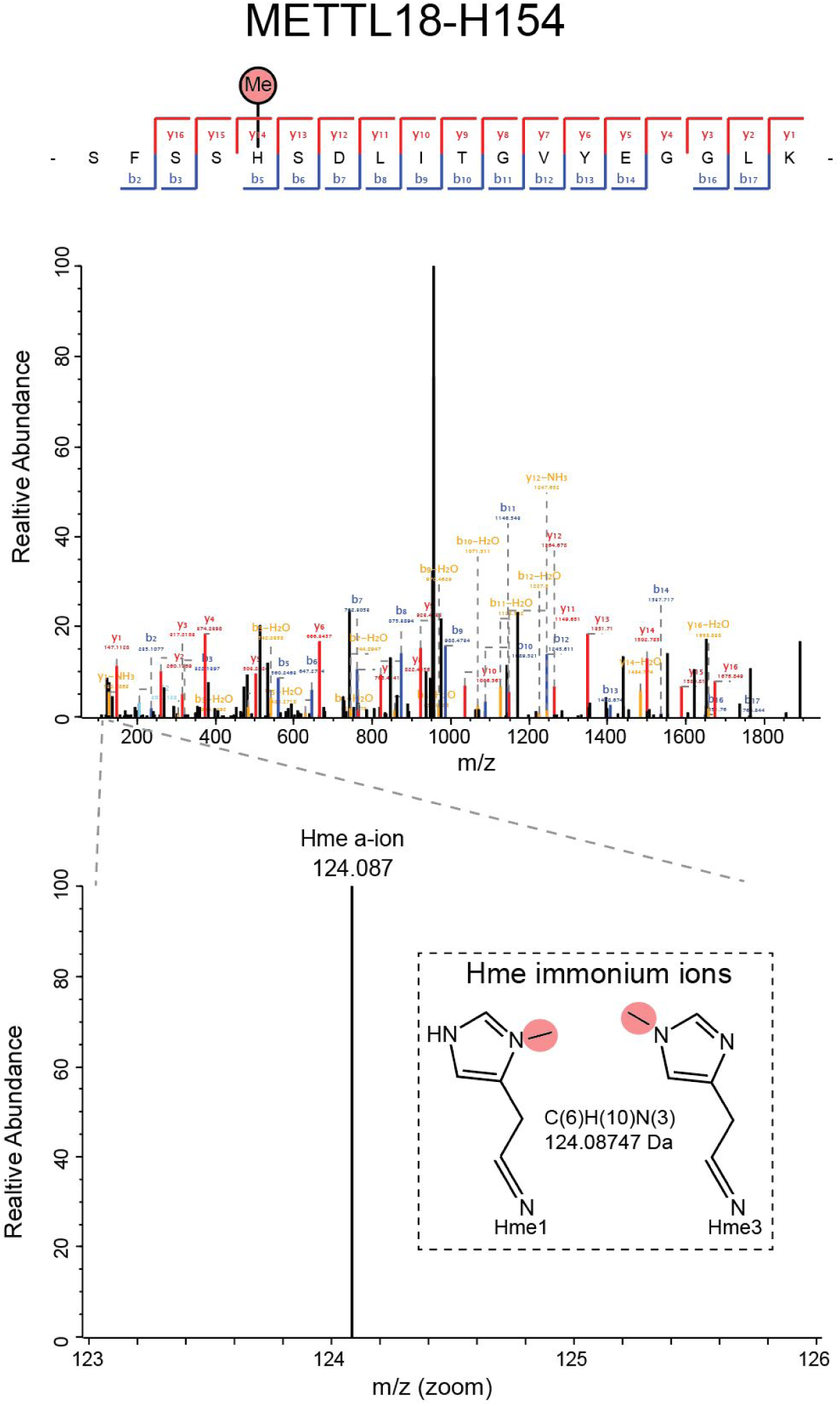
METTL18 is methylated at H154 in HeLa cells. Tandem mass spectrum demonstrating monomethylation of H154 in METTL18 is shown. The peptide corresponds to residues S150-K168 in METTL18 (Uniprot id: O95568).

**Supplementary Figure S5.**
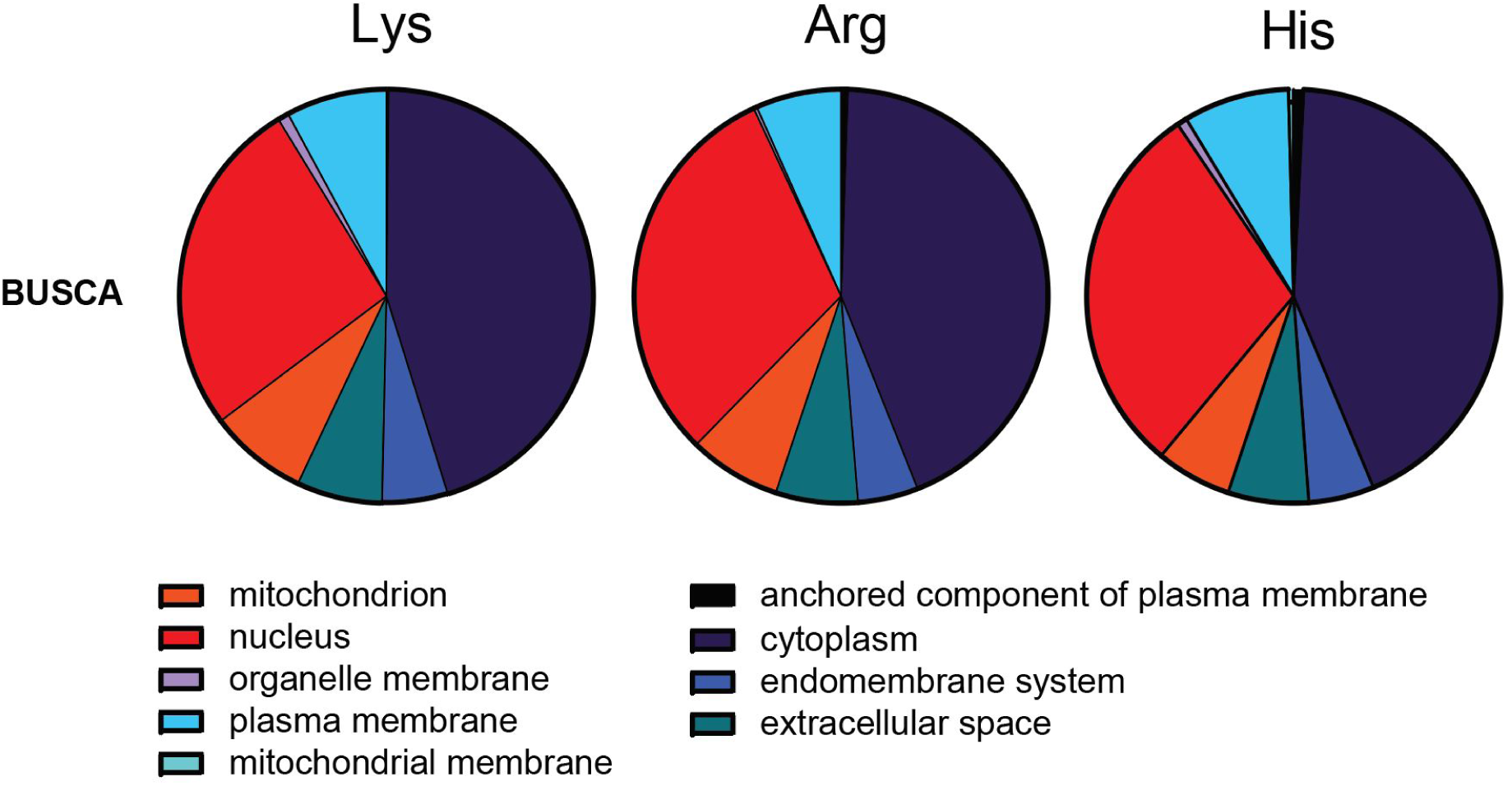
Methylated proteins are distributed throughout the cell. The subcellular localization of proteins methylated in HeLa cells. Pie charts representing the relative predicted localization based on the BUSCA prediction^19^.

**Supplementary Figure S6.**
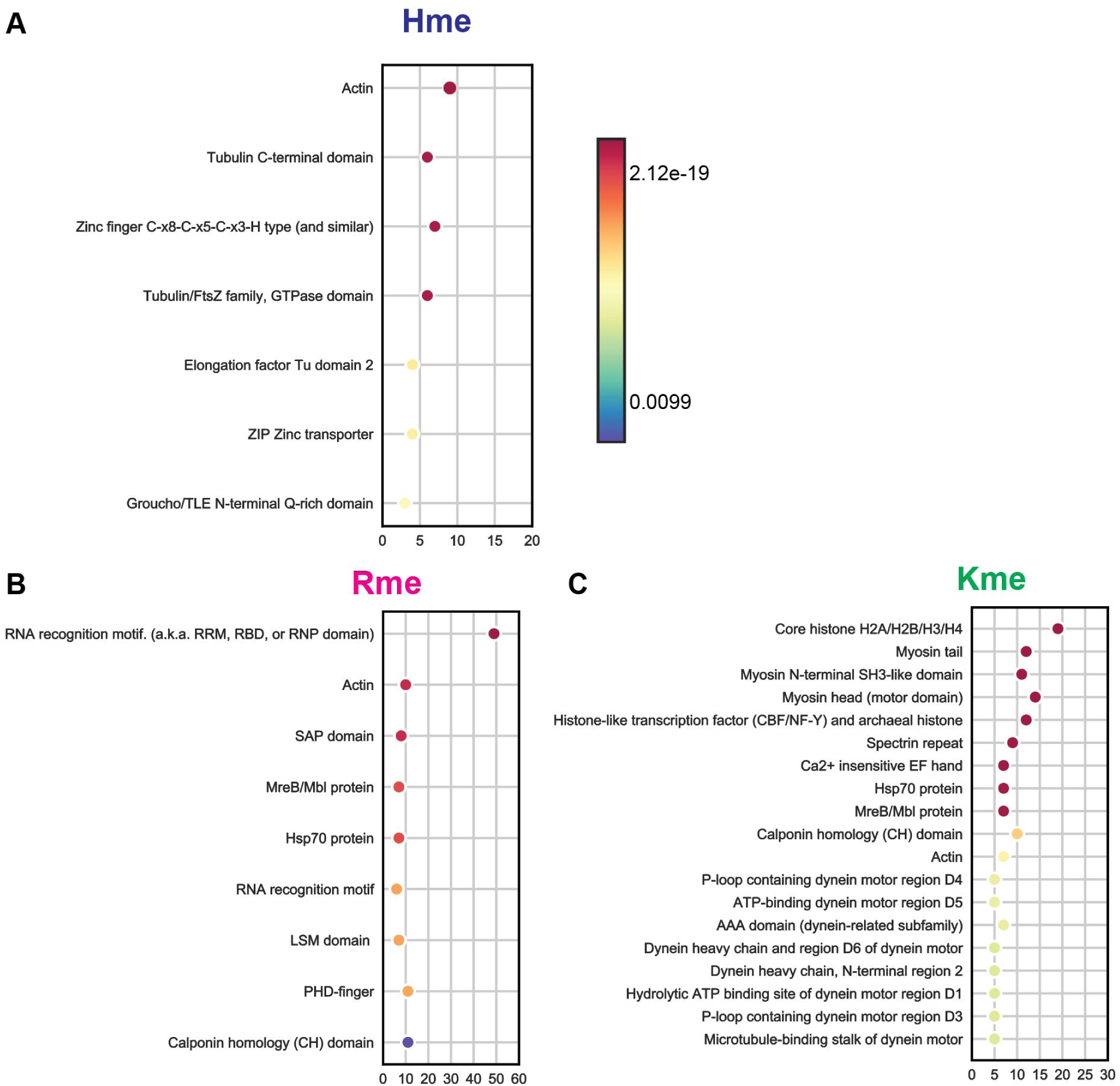
Domain enrichment analysis of proteins containing a methylation site. Functional enrichment analysis performed on methylated proteins based using the Pfam database, categories considered significant (p-value < 0.01) were adjusted using the Benjamini-Hochberg method. The total number of proteins having a significant enrichment in each category is shown on the x-axis.

**Supplementary Figure S7.**
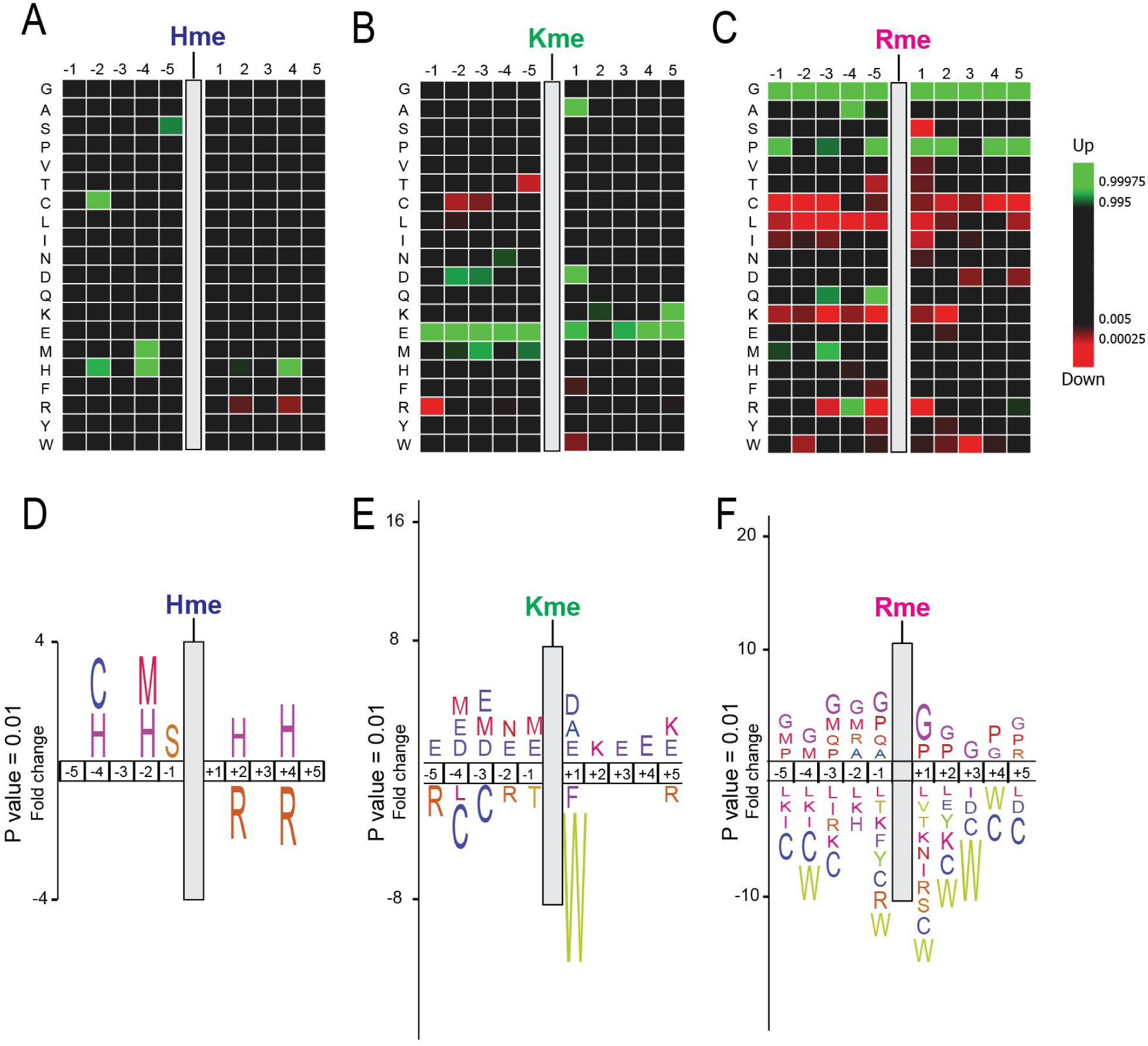
Sequence context of protein methylation events. (**A-C**) Methylome heat maps. Heat maps illustrating over- or under-representation of amino acids in the 5 positions up- and down-stream of identified (A) Hme, (B) Kme, and (C) Rme sites are shown. The enrichment is based on using the *H. Sapiens* precompiled Swiss-Prot composition as reference. (**D-F**) Methylome sequence logos. Logo representations of the heat maps in (A-C) are shown.

## References

1. Johnson, P., Harris, C.I. and Perry, S. V. (1967) 3-Methylhistidine in Actin and Other Muscle Proteins. Biochem. J., 105, 361–370.

2. Huszar, G. and Elzinga, M. (1972) Homologous methylated and nonmethylated histidine peptides in skeletal and cardiac myosins. J. Biol. Chem., 247, 745–753.

3. Kwiatkowski, S. and Drozak, J. (2020) Protein Histidine Methylation. Curr. Protein Pept. Sci., 21, 675–689.

4. Strahl, B. D. and Allis, C. D. (2000) The language of covalent histone modifications. Nature, 403, 41– 45.

5. Kwiatkowski, S., Seliga, A.K., Vertommen, D., Terreri, M., Ishikawa, T., Grabowska, I., Tiebe, M., Teleman, A.A., Jagielski, A.K., Veiga-Da-Cunha, M., et al. (2018) SETD3 protein is the actin-specific histidine N-methyltransferase. Elife, 7, 1–42.

6. Wilkinson, A.W., Diep, J., Dai, S., Liu, S., Ooi, Y.S., Song, D., Li, T.M., Horton, J.R., Zhang, X., Liu, C., et al. (2019) SETD3 is an actin histidine methyltransferase that prevents primary dystocia. Nature, 565, 372–376.

7. Kernstock, S., Davydova, E., Jakobsson, M., Moen, A., Pettersen, S., Mælandsmo, G.M., Egge-Jacobsen, W. and Falnes, P.Ø. (2012) Lysine methylation of VCP by a member of a novel human protein methyltransferase family. Nat. Commun., 3, 1038.

8. Webb, K.J., Zurita-Lopez, C.I., Al-Hadid, Q., Laganowsky, A., Young, B.D., Lipson, R.S., Souda, P., Faull, K.F., Whitelegge, J.P. and Clarke, S.G. (2010) A novel 3-methylhistidine modification of yeast ribosomal protein Rpl3 is dependent upon the YIL110W methyltransferase. J. Biol. Chem., 285, 37598–37606.

9. Davydova, E., Shimazu, T., Schuhmacher, M.K., Jakobsson, M.E., Willemen, H.L.D.M., Liu, T., Moen, A., Ho, A.Y.Y., Schroer, L., Pinto, R., et al. (2021) The methyltransferase METTL9 mediates pervasive 1-methylhistidine modification in mammalian proteomes. Nat. Commun., 10.1038/s41467-020-20670-7.

10. Olsen, J. V. and Mann, M. (2013) Status of Large-scale Analysis of Post-translational Modifications by Mass Spectrometry. Mol. Cell. Proteomics, 12, 3444–3452.

11. Moore, K.E., Carlson, S.M., Camp, N.D., Cheung, P., James, R.G., Chua, K.F., Wolf-yadlin, A. and Gozani, O. (2013) A General Molecular Affinity Strategy for Global Detection and Proteomic Analysis of Lysine Methylation. Mol. Cell, 50, 444–456.

12. Cao, X.J., Arnaudo, A.M. and Garcia, B.A. (2013) Large-scale global identification of protein lysine methylation in vivo. Epigenetics, 8, 477–485.

13. Larsen, S.C., Sylvestersen, K.B., Mund, A., Lyon, D., Mullari, M., Madsen, M. V., Daniel, J.A., Jensen, L.J. and Nielsen, M.L. (2016) Proteome-wide analysis of arginine monomethylation reveals widespread occurrence in human cells. Sci. Signal., 9, 1–15.

14. Devabhaktuni, A., Lin, S., Zhang, L., Swaminathan, K., Gonzalez, C.G., Olsson, N., Pearlman, S.M., Rawson, K. and Elias, J.E. (2019) TagGraph reveals vast protein modification landscapes from large tandem mass spectrometry datasets. Nat. Biotechnol., 37, 469–479.

15. Bekker-Jensen, D.B., Kelstrup, C.D., Batth, T.S., Larsen, S.C., Haldrup, C., Bramsen, J.B., Sørensen, K.D., Høyer, S., Ørntoft, T.F., Andersen, C.L., et al. (2017) An Optimized Shotgun Strategy for the Rapid Generation of Comprehensive Human Proteomes. Cell Syst., 4, 587–599.e4.

16. Deutsch, E.W., Csordas, A., Sun, Z., Jarnuczak, A., Perez-Riverol, Y., Ternent, T., Campbell, D.S., Bernal-Llinares, M., Okuda, S., Kawano, S., et al. (2017) The ProteomeXchange consortium in 2017: Supporting the cultural change in proteomics public data deposition. Nucleic Acids Res., 45, D1100–D1106.

17. Cox, J. and Mann, M. (2008) MaxQuant enables high peptide identification rates, individualized p.p.b.-range mass accuracies and proteome-wide protein quantification. Nat. Biotechnol., 26, 1367–1372.

18. Hornbeck, P. V., Kornhauser, J.M., Tkachev, S., Zhang, B., Skrzypek, E., Murray, B., Latham, V. and Sullivan, M. (2012) PhosphoSitePlus: A comprehensive resource for investigating the structure and function of experimentally determined post-translational modifications in man and mouse. Nucleic Acids Res., 40, 261–270.

19. Orre, L.M., Vesterlund, M., Pan, Y., Arslan, T., Zhu, Y., Fernandez Woodbridge, A., Frings, O., Fredlund, E. and Lehtiö, J. (2019) SubCellBarCode: Proteome-wide Mapping of Protein Localization and Relocalization. Mol. Cell, 73, 166–182.e7.

20. Savojardo, C., Martelli, P.L., Fariselli, P., Profiti, G. and Casadio, R. (2018) BUS CA: An integrative web server to predict subcellular localization of proteins. Nucleic Acids Res., 46, W459–W466.

21. Ochoa, D., Jarnuczak, A.F., Viéitez, C., Gehre, M., Soucheray, M., Mateus, A., Kleefeldt, A.A., Hill, A., Garcia-Alonso, L., Stein, F., et al. (2020) The functional landscape of the human phosphoproteome. Nat. Biotechnol., 38, 365–373.

22. Hunter, S., Apweiler, R., Attwood, T.K., Bairoch, A., Bateman, A., Binns, D., Bork, P., Das, U., Daugherty, L., Duquenne, L., et al. (2009) InterPro: The integrative protein signature database. Nucleic Acids Res., 37, 211–215.

23. Bateman, A., Coin, L., Durbin, R., Finn, R.D., Hollich, V., Griffiths-Jones, S., Khanna, A., Marshall, M., Moxon, S., Sonnhammer, E.L.L., et al. (2004) The Pfam protein families database. Nucleic Acids Res., 32, 138–141.

24. Szklarczyk, D., Gable, A.L., Lyon, D., Junge, A., Wyder, S., Huerta-Cepas, J., Simonovic, M., Doncheva, N.T., Morris, J.H., Bork, P., et al. (2019) STRING v11: Protein-protein association networks with increased coverage, supporting functional discovery in genome-wide experimental datasets. Nucleic Acids Res., 47, D607–D613.

25. Colaert, N., Helsens, K., Martens, L., Vandekerckhove, J. and Gevaert, K. (2009) Improved visualization of protein consensus sequences by iceLogo. Nat. Methods, 6, 786–787.

26. Kelstrup, C.D., Jersie-Christensen, R.R., Batth, T.S., Arrey, T.N., Kuehn, A., Kellmann, M. and Olsen, J. V. (2014) Rapid and deep proteomes by faster sequencing on a benchtop quadrupole ultra-high-field orbitrap mass spectrometer. J. Proteome Res., 13, 6187–6195.

27. Jakobsson, M.E., Małecki, J.M., Halabelian, L., Nilges, B.S., Pinto, R., Kudithipudi, S., Munk, S., Davydova, E., Zuhairi, F.R., Arrowsmith, C.H., et al. (2018) The dual methyltransferase METTL13 targets N terminus and Lys55 of eEF1A and modulates codon-specific translation rates. Nat. Commun., 9, 3411.

28. Tyanova, S., Temu, T., Sinitcyn, P., Carlson, A., Hein, M.Y., Geiger, T., Mann, M. and Cox, J. (2016) The Perseus computational platform for comprehensive analysis of (prote)omics data. Nat. Methods, 13, 731–740.

29. Potel, C.M., Lin, M.H., Prust, N., Van Den Toorn, H.W.P., Heck, A.J.R. and Lemeer, S. (2019) Gaining Confidence in the Elusive Histidine Phosphoproteome. Anal. Chem., 91, 5542–5547.

30. Trelle, M.B. and Jensen, O.N. (2008) Utility of immonium ions for assignment of ε-N-acetyllysine-containing peptides by tandem mass spectrometry. Anal. Chem., 80, 3422–3430.

31. Steen, H. and Mann, M. (2004) The ABC’s (and XYZ’s) of peptide sequencing. Nat. Rev. Mol. Cell Biol., 5, 699–711.

32. Ly, T. and Lamond, A.I. (2017) New Apex in Proteome Analysis. Cell Syst., 4, 581–582.

33. Ning, Z., Star, A.T., Mierzwa, A., Lanouette, S., Mayne, J., Couture, J.F. and Figeys, D. (2016) A charge-suppressing strategy for probing protein methylation. Chem. Commun., 52, 5474–5477.

34. Bantscheff, M., Schirle, M., Sweetman, G., Rick, J. and Kuster, B. (2007) Quantitative mass spectrometry in proteomics: A critical review. Anal. Bioanal. Chem., 389, 1017–1031.

35. Falnes, P.Ø., Jakobsson, M.E., Davydova, E., Ho, A. and Małecki, J. (2016) Protein lysine methylation by seven-β-strand methyltransferases. Biochem. J., 473, 1995–2009.

36. Turner, B.M. (2002) Cellular memory and the histone code. Cell, 111, 285–291.

37. Jakobsson, M.E., Małecki, J., Nilges, B.S., Moen, A., Leidel, S.A. and Falnes, P.Ø. (2017) Methylation of human eukaryotic elongation factor alpha (eEF1A) by a member of a novel protein lysine methyltransferase family modulates mRNA translation. Nucleic Acids Res., 45, 8239–8254.

38. Jakobsson, M.E., Małecki, J. and Falnes, P.Ø. (2018) Regulation of eukaryotic elongation factor 1 alpha (eEF1A) by dynamic lysine methylation. RNA Biol., 6286, 01–11.

39. Jakobsson, M.E., Moen, A., Bousset, L., Egge-Jacobsen, W., Kernstock, S., Melki, R. and Falnes, P. (2013) Identification and characterization of a novel human methyltransferase modulating Hsp70 protein function through lysine methylation. J. Biol. Chem., 288, 27752–27763.

40. Jakobsson, M.E., Moen, A. and Falnes, P.O. (2016) Correspondence: On the enzymology and significance of HSPA1 lysine methylation. Nat Commun, 7, 11464.

41. Szewczyk, M.M., Ishikawa, Y., Organ, S., Sakai, N., Li, F., Halabelian, L., Ackloo, S., Couzens, A.L., Eram, M., Dilworth, D., et al. (2020) Pharmacological inhibition of PRMT7 links arginine monomethylation to the cellular stress response. Nat. Commun., 11, 1–15.

42. Sonnhammer, E.L.L., Eddy, S.R., Birney, E., Bateman, A. and Durbin, R. (1998) Pfam: Multiple sequence alignments and HMM-profiles of protein domains. Nucleic Acids Res., 26, 320–322.

43. Sabbattini, P., Sjoberg, M., Nikic, S., Frangini, A., Holmqvist, P.H., Kunowska, N., Carroll, T., Brookes, E., Arthur, S.J., Pombo, A., et al. (2014) An H3K9/S10 methyl-phospho switch modulates Polycomb and Pol II binding at repressed genes during differentiation. Mol. Biol. Cell, 25, 904– 915.

44. Estéve, P.O., Chang, Y., Samaranayake, M., Upadhyay, A.K., Horton, J.R., Feehery, G.R., Cheng, X. and Pradhan, S. (2011) A methylation and phosphorylation switch between an adjacent lysine and serine determines human DNMT1 stability. Nat. Struct. Mol. Biol., 18, 42–49.

45. Liu, X., Wu, C., Li, C. and Boerwinkle, E. (2016) dbNSFP v3.0: A One-Stop Database of Functional Predictions and Annotations for Human Nonsynonymous and Splice-Site SNVs. Hum. Mutat., 37, 235–241.

46. Landrum, M.J., Lee, J.M., Riley, G.R., Jang, W., Rubinstein, W.S., Church, D.M. and Maglott, D.R. (2014) ClinVar: Public archive of relationships among sequence variation and human phenotype. Nucleic Acids Res., 42, 980–985.

47. Garcia-Angarita, N., Kirschner, J., Heiliger, M., Thirion, C., Walter, M.C., Schnittfeld-Acarlioglu, S., Albrecht, M., Müller, K., Wieczorek, D., Lochmüller, H., et al. (2009) Severe nemaline myopathy associated with consecutive mutations E74D and H75Y on a single ACTA1 allele. Neuromuscul. Disord., 19, 481–484.

48. Dominguez, R. and Holmes, K.C. (2011) Actin structure and function. Annu. Rev. Biophys., 40, 169–186.

49. Essletzbichler, P., Konopka, T., Santoro, F., Chen, D., Gapp, B. V., Kralovics, R., Brummelkamp, T.R., Nijman, S.M.B. and Bürckstümmer, T. (2014) Megabase-scale deletion using CRISPR/Cas9 to generate a fully haploid human cell line. Genome Res., 24, 2059–2065.

50. Olsen, J. V, Vermeulen, M., Santamaria, A., Kumar, C., Miller, M.L., Jensen, L.J., Gnad, F., Cox, J., Jensen, T.S., Nigg, E.A., et al. (2010) Quantitative phosphoproteomics reveals widespread full phosphorylation site occupancy during mitosis. Sci. Signal., 3, ra3.

51. Olsen, J. V, Blagoev, B., Gnad, F., Macek, B., Kumar, C., Mortensen, P. and Mann, M. (2006) Global, In Vivo, and Site-Specific Phosphorylation Dynamics in Signaling Networks. Cell, 127, 635–648.

52. Choudhary, C., Kumar, C., Gnad, F., Nielsen, M.L., Rehman, M., Walther, T.C., Olsen, J. V and Mann, M. (2009) Lysine acetylation targets protein complexes and co-regulates major cellular functions. Science (80-.)., 325, 834–40.

53. Akimov, V., Barrio-Hernandez, I., Hansen, S.V.F., Hallenborg, P., Pedersen, A.K., Bekker-Jensen, D.B., Puglia, M., Christensen, S.D.K., Vanselow, J.T., Nielsen, M.M., et al. (2018) Ubisite approach for comprehensive mapping of lysine and n-terminal ubiquitination sites. Nat. Struct. Mol. Biol., 25.

54. Hendriks, I.A., Lyon, D., Young, C., Jensen, L.J., Vertegaal, A.C.O. and Nielsen, M.L. (2017) Site-specific mapping of the human SUMO proteome reveals co-modification with phosphorylation. Nat. Struct. Mol. Biol., 24, 325–336.

55. Gray, A., Bradbury, A.R.M., Knappik, A., Plückthun, A., Borrebaeck, C.A.K. and Dübel, S. (2020) Animal-free alternatives and the antibody iceberg. Nat. Biotechnol., 38, 1234–1239.

56. Kehoe, J.W., Velappan, N., Walbolt, M., Rasmussen, J., King, D., Lou, J., Knopp, K., Pavlik, P., Marks, J.D., Bertozzi, C.R., et al. (2006) Using Phage Display to Select Antibodies Recognizing Post-translational Modifications Independently of Sequence Context. Mol. Cell. Proteomics, 5, 2350–2363.

57. Gayatri, S., Cowles, M.W., Vemulapalli, V., Cheng, D., Sun, Z.W. and Bedford, M.T. (2016) Using oriented peptide array libraries to evaluate methylarginine-specific antibodies and arginine methyltransferase substrate motifs. Sci. Rep., 6, 1–8.

58. Bock, I., Kudithipudi, S., Tamas, R., Kungulovski, G., Dhayalan, A. and Jeltsch, A. (2011) Application of Celluspots peptide arrays for the analysis of the binding specificity of epigenetic reading domains to modified histone tails. BMC Biochem., 12, 48.

59. Varland, S., Vandekerckhove, J. and Drazic, A. (2019) Actin Post-translational Modifications: The Cinderella of Cytoskeletal Control. Trends Biochem. Sci., 44, 502–516.

60. Aksnes, H., Ree, R. and Arnesen, T. (2019) Co-translational, Post-translational, and Non-catalytic Roles of N-Terminal Acetyltransferases. Mol. Cell, 73, 1097–1114.

61. Petrossian, T.C. and Clarke, S.G. (2011) Uncovering the human methyltransferasome. Mol. Cell. Proteomics, 10, M110.000976.

62. Meyer, K. and Selbach, M. (2020) Peptide-based interaction proteomics. Mol. Cell. Proteomics, 19, 1070–1075.

63. Peterson, A.C., Russell, J.D., Bailey, D.J., Westphall, M.S. and Coon, J.J. (2012) Parallel reaction monitoring for high resolution and high mass accuracy quantitative, targeted proteomics. Mol. Cell. Proteomics, 11, 1475–1488.

64. Perez-Riverol, Y., Csordas, A., Bai, J., Bernal-Llinares, M., Hewapathirana, S., Kundu, D.J., Inuganti, A., Griss, J., Mayer, G., Eisenacher, M., et al. (2019) The PRIDE database and related tools and resources in 2019: Improving support for quantification data. Nucleic Acids Res., 47, D442–D450.

